# Dissociation of movement and outcome representations in metacognition of agency

**DOI:** 10.1101/2024.10.14.618169

**Authors:** Angeliki Charalampaki, Heiko Maurer, Lisa K. Maurer, Hermann Müller, Elisa Filevich

## Abstract

We studied the role of movement and outcome information in forming metacognitive representations of agency. Participants completed a goal-oriented task, a virtual version of a ball-throwing game. In two conditions, we manipulated either the visual representation of the throwing movement or its distal outcome (the resulting ball flight/trajectory). We measured participants’ accuracy in a discrimination agency task, as well as confidence in their responses and tested for differences in the electrophysiological (EEG) signal using linear mixed effect models. We found no mean differences between participants’ metacognitive efficiency between conditions, but we also found that metacognitive sensitivity did not correlate between the two conditions, suggesting a dissociation in their underlying mechanisms. Furthermore, exploratory analyses pointed toward a difference in the EEG signal between the two conditions. Taken together, our results suggest that while movement and outcome information contribute equally to participants’ sense of agency, they may do so through distinct underlying processes.

## Introduction

The subjective experience that we are the author of our intentional actions is referred to as the sense of agency. This is an umbrella term that can refer to many potentially distinct cognitive processes. In fact, previous work has pointed at this breadth of the scope in the term and aimed to distinguish different operational definitions of agency (sense of intentional causing an action vs. sense of initiating an action vs. sense of control: Pacherie, 2007; pre-reflective vs. reflective components of agency: Gallagher, 2012; feeling of agency vs. judgment of agency vs meta-representational level: Synofzik et al., 2008; Christensen & Grünbaum, 2018; agency as an ability vs. phenomenal feel and bodily vs. external agency: Christensen & Grünbaum, 2017; Wen, 2019 just to name a few). The different aspects encompassed by the broad concept of the sense of agency will arguably have different underlying mechanisms and must be studied experimentally in different ways. But in previous studies, the details of the operational definitions are frequently ignored, possibly explaining the lack of consensus on the brain regions linked to this subjective experience (Charalampaki et al., 2022). In particular, there is an ongoing discussion on the similarities and differences between bodily and external operational definitions of agency (Christensen & Grünbaum, 2017; Kalckert & Ehrsson, 2012; Wen, 2019). The difference lies in whether the definition used in a study of agency over an action (“I threw the ball in the basket”) is strictly limited by the boundaries of our body (the movement: “I moved my arm to throw”) or the definition is broad enough to also include agency over the consequences of our actions that go beyond the boundaries of our body (the outcome: “The ball in the basket”). In what follows, we will first discuss the most common model used to explain the human sense of agency: the comparator model (Frith et al., 2000), which is often used interchangeably to describe both bodily and external sense of agency. While there is evidence that agency over the movement and agency over the outcome is diminished by similar types of experimental manipulations, conflicting results exist on their relative importance in our sense of agency, which challenges the notion that we can treat them interchangeably. We will then conclude with how our study can inform this discussion by borrowing methods from the metacognitive literature.

According to the comparator model, whenever we make an intentional action, a copy of the motor (efferent) commands issued to make the necessary movements is used to make predictions about the consequences of the movement. These predictions are then compared with the actual afferent sensory feedback (associated with our moving limbs). When these two match, we typically experience authorship of the movement; when they do not, we experience a loss of agency. While the comparator model was originally used to explain agency over our bodily movements (Frith et al., 2000), it is often extended to explain the agency we experience over the external consequences accompanying our actions (Haggard, 2017). That is, it is usually assumed that the comparator can also make comparisons between predictions and feedback for the sensory consequences that go beyond our bodies. However, this assumption has been challenged. In particular, because current experimental designs have either failed to or provided mixed results regarding the involvement of efference copies in predicting the sensory consequences of outcomes beyond the body (Christensen & Grünbaum, 2018a; Dogge et al., 2019). Further, participants can experience agency even for actions carried out by another person (Tieri et al., 2015; Wegner et al., 2004). Meaning that agency can be experienced in the absence of an efference signal (but see Kilteni et al., 2021 for the importance of efference copies in the sensory attenuation). Therefore, it is important to question the suitability of the comparator model to explain the sense of agency over external consequences of movement. On the one hand, one series of studies shows that subjective reports on participants’ sense of agency is affected both by manipulations of visual representations of a movement or its outcome. Participants reported experiencing a diminished sense of agency over an action when they fail to achieve a specific goal (van der Weiden et al., 2013), when the outcome was different than their predictions (Sato & Yasuda, 2005), or when the predicted outcome was delayed (Farrer et al., 2013; Spengler et al., 2009). Similarly, participants reported a lower sense of agency over a movement they made when the visual feedback of the movement they performed was manipulated spatially (Farrer et al., 2003; Padrao et al., 2016; van den Bos & Jeannerod, 2002) or temporally (Farrer et al., 2003; Krugwasser et al., 2019).

On the other hand, another line of work has aimed at directly comparing the relative precision of agency representations based on movement and outcome prediction violations, with conflicting results. While some studies found that participants were more sensitive to violations of outcome predictions as compared to violations of bodily predictions, other studies have found the opposite. For example, David et al. (2016) introduced a sensorimotor mismatch by temporally delaying the visual feedback, either at the point of movement initiation or at the onset of the movement outcome. They found that participants’ agency ratings decreased most sharply when the timing of the outcome was manipulated. Similarly, Wen et al. (2015) had participants click on a keyboard to control the position of a moving dot, aiming to make rapid movements to reach a visual target. The visual feedback was manipulated by introducing a temporal lag, and the level of control on the movement was manipulated by ignoring some of the participants’ commands. The latter manipulation was meant to introduce a sensorimotor mismatch so as to affect whether participants reached their goal, sometimes improving their success rate. Participants’ agency reports reflected whether they succeeded with their goal despite some of their commands being ignored, albeit only under some conditions, where the delay introduced was long. Finally, Oishi et al. (2018) extended these findings by showing that reaching a goal can retrospectively influence participants’ agency, even when it is independent of their performance. In their study, participants had to move a cursor (in the presence or absence of temporal delays) toward a changing color target, with the goal of reaching the target while it had a specific color – which was out of the participants’ control. Participants then rated the level of control over the movement of the cursor (Oishi et al., 2018).

The studies summarized above point toward outcome prediction overriding movement prediction violations, suggesting in turn that the representation of outcome predictions might affect participants’ agency more strongly than representations of movement predictions. But, as mentioned above, other studies directly contradict this conclusion, by showing that participants are more sensitive to violations of motor predictions as compared to violations of outcome predictions. For example, Metcalfe and Green (2007) had participants move a cursor toward downward-moving visual targets and, on some trials, manipulated the movement feedback by introducing “turbulence” on the cursor’s movement. They also manipulated the outcome by introducing false outcome hits (namely the target appeared as being "magically" touched). Without turbulence, participants’ agency ratings (level of control) matched the proportion of targets they hit. In other words, in the absence of movement prediction violations, the more targets they hit, the higher they rated the level of control. However, when there were either movement manipulations or outcome manipulations, participants’ agency ratings more closely followed the relation between their motor act and the visual feedback of it, irrespective of how many targets it appeared as they had touched. In a similar study, Metcalfe et al. (2013) extended their findings by manipulating both the motor aspect of the action (turbulence) and the probability of a target being exploded upon touch. They showed that while both types of violations negatively affected participants’ agency, violations of the movement predictions had a more notable effect on participants’ ratings of control, and this was independent of participants’ performance, measured as the number of targets participants hit.

As a whole, this body of literature demonstrates that both bodily and external prediction violations, probed with manipulations of the feedback on either the movement or its outcome, can affect explicit reports. Importantly however, the different results also emphasize the importance —and challenges— of differentiating between different experimental manipulations in the study of agency. Beyond the conceptual differences driven by different experimental operationalizations of the sense of agency, there are two methodological aspects that limit the interpretability of the findings: All of the experiments summarized above measure participants’ sense of agency using subjective reports, which are prone to biases and this could explain the apparent diversity of the findings regarding the relative importance of movement and outcome predictions in our subjective experience of agency. Also, some of the studies compared participants sensitivity in detecting prediction violations of movement and outcome using manipulations that differ substantially between the two conditions. For example, manipulating the movement feedback with turbulence vs. manipulating the outcome feedback by controlling the probability of an outcome occurring (Metcalfe et al., 2013). This challenges the comparison of the precision of these two sources of information as we do not know how these two different levels of action description relate to one another. To deal with these complications, while studying agency with explicit reports, one can instead measure the precision of explicit representations of agency using tools borrowed from research in metacognition (Fleming & Lau, 2014). That is, one can measure metacognitive representations of the agency judgments that lie on one representational level above, allowing for a comparison of their relative precision. This was the approach we chose.

In this study, participants made movements in a deterministic, goal-oriented motor task in which we could tightly manipulate both the visual representations of the movement and the outcome. The motor task was a semi-virtual version of the "Skittles" game (Müller & Sternad 2004; Fig. 1). In the Skittles game, in which a ball is attached to a pole with a rope, the aim is to grab and throw the ball in such a way to hit the target placed behind the pole. In this task, we could manipulate both the visual representations of the movement and the outcome. On each trial, participants completed two consecutive motor actions (grab and throw the ball to hit the target); they selected which of the two actions they felt were more in control (only one matched with their action) and then rated their confidence in that decision. We hypothesized that representations of movement and outcome differentially affect participants’ metacognitive representations of their sense of agency, and that this would also be reflected as behavioral differences. We also explored possible differences at the neural level in the underlying electroencephalography (EEG) data.

**Fig. 1.**
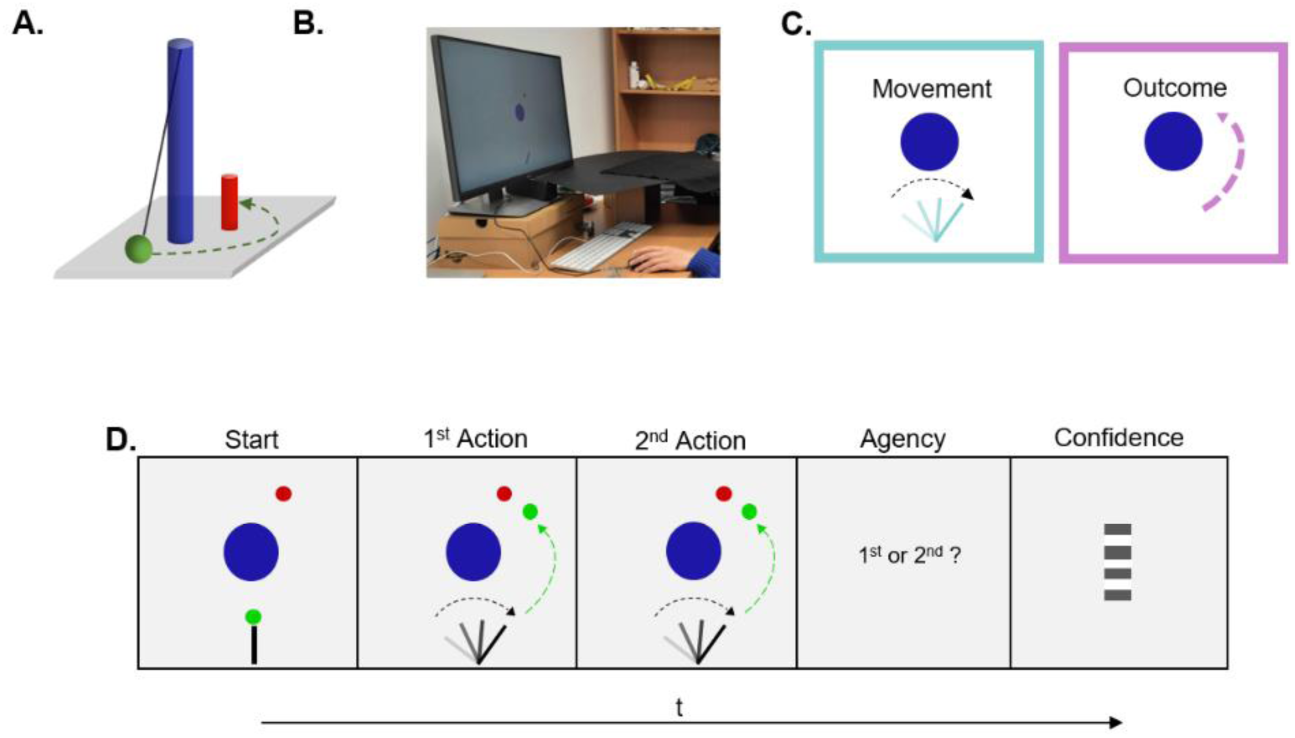
A. The Skittles game. The aim of the game is to swing the ball (green) attached to the pole (blue) with a rope to hit the target (red). **B. Experimental set-up.** Participants sat in front of the computer, placed their right hand on the custom-made lever (which was hidden below an occluder), and used a computer mouse with their left hand to register their responses. We recorded EEG signal from 62 active electrodes placed on an elastic head cap. **C. Experimental conditions.** We manipulated the online visual feedback participants received on the screen in two conditions. In the *Movement* condition, in only one of the two intervals, we manipulated the position of the virtual lever, which moved synchronously with the physical one but deviated a few degrees to the left or right. In the *Outcome* condition, in only one of the two intervals, we manipulated the ball trajectory. The ball flew in an alternative trajectory with the same distal outcome (hit/miss the target) as the participants’ actual trajectory. **D. Experimental paradigm.** In the main experiment, participants threw the ball two consecutive times and then selected in which of the two they felt more in control. Finally, participants rated how confident they were about their decision.

## Methods

We assessed whether violations of visual representations of a movement and its consequences on the environment equivalently affect participants’ sense of agency using behavioral and electrophysiological data (EEG). The study was pre-registered (https://osf.io/sjurk), and we report any deviations from the pre-registered plan. The ethics committee of the Institute of Psychology of the Humboldt-Universität zu Berlin approved the experimental procedures (2017-17), which were in line with the Declaration of Helsinki.

### Participants

Our pre-registered plan was to collect 40 clean datasets from participants who passed the inclusion criteria: no neurological or psychiatric history, normal or corrected-to-normal vision, right-handedness, and proficiency in English. The sample size was determined based on a previous study in which the Skittles task was embedded in a motor metacognitive conducted experimental design (Arbuzova et al., 2021). To that extent, we collected data from 52 participants. From these datasets, we excluded five participants with very low discrimination accuracy (< 60%) in at least one of the two conditions. We only included participants who correctly selected the interval in which they were in control (the online feedback matched their action), 60% or more of the total number of trials belonging to each condition. In deviating from our pre-registered plan, we further excluded seven datasets in which discrimination performance was especially variable throughout the experimental session, determined by visually inspecting the adaptively changed manipulation magnitude (see the description of the staircasing procedure below) prior to analyzing the data. This resulted in a final sample of 40 participants (mean age = 26.60 years ± 5.60, 25 female, 15 male, 0 diverse, mean handedness score = 90.60 ± 14.50 [Oldfield, 1971]). Due to technical issues the EEG signal from two participants was not recorded, but we nevertheless used these participants’ data in the behavioral analysis. Participants signed a written informed consent, and for their time, they were compensated with 8 €/hour or course credits.

### Apparatus and stimuli

For the motor task, we adapted the virtual version of the "Skittles" game, first described by Müller and Sternad (2004). In the Skittles game, a ball is attached to a pole with a rope, and a target –the Skittle– is placed behind the pole (Fig. 1. Panel A.). The goal of the game is to knock down the target by swinging the ball around the pole. In the virtual version of the Skittles game, participants had a top-down view of the scene: The central pole was placed in the middle of the scene (blue circle, position (x, y) = (0,0) m) with a radius = 0.25 m); the ball originally appeared hanging next to the pole (green circle, radius = 0.05 m) and the target was, when viewed from above, top-right of the pole (red circle, position (x,y) = (0.20, 0.70) m, radius = 0.05). The massless rope (spring constant k = 1 N/m) was not visible on the scene, but was taken into consideration when estimating the ball trajectory. Participants grabbed, moved, and released the ball using a custom-made lever. The lever rotated on the end nearest to the participants, so that they could swing their arm horizontally around the elbow point. The angle of their hand was recorded online with the aid of a goniometer (Novotechnik, Stuttgart, Germany, RFC4800 Model 600, 12-bit resolution, 0.1° precision), and all the data from the lever were recorded with a sampling rate of 1 kHz using a Labjack T7 data acquisition device (LabJack Corp., Lakewood, CO). This allowed us to depict the virtual lever in the lower middle part of the scene ((x, y) = (0,-1.5) m) as a black rectangular (0.4 m length, 0.04 width) that pivoted just like the physical lever. Participants grabbed the virtual ball by placing their index finger on the touch switch on the tip of the lever and released it by lifting their finger. The Skittles game is deterministic. That is, the ball trajectory can be determined at the point of ball release by the angle of the lever and the velocity of its tip (Müller & Sternad, 2004; Sternad et al., 2011). Once participants lifted their finger, the ball disappeared for approximately 200 ms (M = 202.36 ± 0.21) to avoid a jump in the ball position when the ball trajectory was manipulated (see below). Therefore, we could display the ball trajectory in real-time with a maximum possible flight duration of 2 s. Because on each trial of the metacognitive task (see below) participants would compare two actions, we aimed to match the goal of each one participants’ action (Charalampaki et al., 2023). We therefore emphasized to the participants to always aim to hit the target, and we positioned the target slightly on the right of the pole to increase the chances that participants hit it. Each motor task started with only the central pole displayed, and with a pseudo-randomized delay (0.40 to 1.00 s), the lever, the ball, and the target appeared on the screen. We programmed the task in MATLAB (R2016b, MathWorks, Natick, MA) and Psychtoolbox-3 (Brainard, 1997; Kleiner et al., 2007; Pelli, 1997). Throughout the experiment, participants had no visual access to either their arm or the real lever (an opaque piece of cloth was held above their arm, without interfering with the movement), so they had to rely on the visual feedback they received on an LCD monitor (2560 × 1440 pixels, 0.61 × 0.345 m, refresh rate 60 Hz) located approximately 0.6 m in front of them (Fig. 1. Panel B.).

### Procedure

On each trial, participants performed the motor task two consecutive times. The online visual feedback they received matched their action in only one of the two intervals (Fig. 1. Panel C.). In the other, the visual feedback was manipulated in one of two possible ways: either the position of the lever (*Movement* condition) or the trajectory of the ball (*Outcome* condition) did not match their own. Two previous studies used the semi-virtual Skittles game and found that participants had accurate representations of both movement parameters –the velocity of the movement and the hand’s angle position at the point of ball release– but also the ball flight trajectory and the distal outcome (Arbuzova et al., 2021; Charalampaki et al., 2023). We used a visuospatial manipulation to introduce violations of the predictions of the movement and outcome for two reasons. First, because temporal delay could affect participants ability to monitor their action rather than simply introducing a mismatch between the prediction and the feedback of their action (for an extended discussion on the topic see Wen, 2019; but also Osumi et al., 2019) and second, to prevent participants from recognizing the visually delayed feedback as a delayed version of their action (Farrer et al., 2008). In the *Movement* condition, we only manipulated the angle of the lever (see details on the *Online staircases section*). Therefore, the virtual lever always moved synchronously with the physical one but had a different angle, and the ball flew according to the participant’s actual throw (based on the angle of participants’ real hand and velocity at the point of ball release). In the *Outcome* condition, the virtual lever precisely matched the participant’s hand’s movement. We manipulated only the ball trajectory, ensuring that it would always match the distal outcome of the participants’ real throw. At the end of the second action, participants discriminated in a two-interval forced choice task, in which of the two intervals they felt more in control (Fig. 1. Panel D.). To do so, they could press the left or right computer mouse buttons with their left hand to report the first or second interval, respectively. Following the discrimination response, participants rated how confident they were that the interval they selected was the one they had complete control over the action seen on the screen using a continuous virtual scale from "Very unsure" to "Very confident". Participants moved up and down a computer mouse with their left hand, which controlled the position of a bar that appeared on a pseudo-random position over the virtual scale on each trial, and then pressed either button on the mouse to register their confidence. During this point of the trial, participants could alternatively press the spacebar on a computer keyboard to mark the trial for exclusion (e.g., they pushed the wrong button in the discrimination step).

To familiarize participants with the virtual version of the task, participants first completed eight trials of the motor task in which the visual feedback always matched their action. After that, they did 40 training trials, half of which were *Movement* and half *Outcome* trials presented in pseudo-randomised interleaved order. In these training trials, the trial ended after the discrimination response. Finally, participants completed 16 trials in which they also rated their confidence in their decision to become familiar with the steps of the main experiment. The main experiment consisted of 504, split into 9 blocks, with breaks in between. Each block consisted of 56 pseudo-randomized and interleaved *Movement* and *Outcome* condition trials. The whole experiment, including EEG electrodes elastic cap preparation, took approximately three hours.

### Online staircases

To keep participants’ discrimination accuracy around 71% (Levitt, 1971), we used a separate online staircase (2-down, 1-up) for each condition. In both cases, we staircased the value |Δv| that we used to manipulate the different aspects of the motor task for each of the two conditions. Specifically, on *Outcome* trials we manipulated the trajectory of the ball by changing the velocity of the real movement in |Δv|, in order to create a manipulated ball flight trajectory. We pseudorandomized a parameter to determine whether we added or subtracted |Δv|, in order to present a manipulated trajectory to the right or left of the real one, respectively. We deviated from this process if the alternative ball trajectory ended in a throw with a distal outcome (hit/miss the target) that did not match participants’ throw. In these instances, we first multiplied the |Δv| with the opposite sign, and if this did not work either, we would use the nearest possible value that would. If none of these steps worked, we used the original staircase value and marked the trial for exclusion. On *Movement trials,* we also staircased |Δv|, but first multiplied it by the time difference Δt between two screen refresh frames in order to obtain the lever angle difference corresponding to the staircased |Δv|. Importantly, on Outcome trials, in both intervals, the lever position corresponded to the participant’s hand position, and on Movement trials, the ball flight trajectory in both intervals corresponded to the trajectory based on the participant’s hand position at the point of ball release. The starting value of both staircases was 1, the step size was 0.15 in the training session and 0.10 in the main experiment, and the minimum possible value of the staircase was 0.21 for the training and 0.01 in the main experiment. The larger values used in the training staircases were meant to reach faster those values that would stabilize participants’ performance. We then used the last values of each staircase of the training session as the starting values for the staircases used in the main experiment.

### Metacognitive representations of agency

This design allowed us to quantify how well people can inspect their agency judgments. That is, we could use an estimate of metacognitive efficiency (Mratio; Maniscalco & Lau, 2012), which reflects the relationship between participants’ discrimination accuracy and confidence rating. Mratio aims to quantify the proportion of information available for confidence ratings, relative to that available in the discrimination decisions. Thus, as a unit-free measure, it allows for the comparison between conditions without being contaminated by potential biases in subjective reports. Participants with high Mratio typically rate with high confidence trials in which they correctly identified in which of two intervals they had more control of the perceived action; and rate with low confidence those trials in which they instead identified the interval in which they had less control of the perceived action. In contrast, confidence ratings from participants with low Mratio typically cannot differentiate between the trials in which they selected the correct interval from those they did not.

## Data analysis

### Data exclusion

In keeping with our pre-registered plan, we excluded from both the behavioral and EEG data those trials in which the discrimination reaction time was above 8 s (mean excluded trials ± SD: 1 ± 2), trials marked for exclusion by the participants (mean ± SD: 4 ± 6), and those trials in which the intervals differed in terms of the distal outcome (mean ± SD of trials participants missed hitting the target in either of the intervals: 63 ± 55 and mean trials in which participants threw the ball towards the left of the pole in at least one interval: 5 ± 11). Overall, we included 405 ± 70 trials (Movement trials: 198 ± 38 and Outcome trials: 207 ± 33).

In addition to our pre-registered exclusion criteria, we excluded the (behavioral and EEG) data from the first blocks from those participants for whom the staircases were especially variable at the start of the experiment. Specifically, we excluded the first block from eight participants, the first two blocks from four participants, and the first four blocks from two participants. Data exclusion took place prior to any data analysis.

### Behavioral data

To estimate the precision of participants’ metacognition of agency, we measured their metacognitive efficiency (Mratio; Maniscalco & Lau, 2012) using the maximum likelihood estimation method as implemented in the *“metaSDT”* R package (Craddock, 2021). For this analysis, we had to discretize the continuous confidence ratings, and we chose to do so using six equidistant bins. Before discretizing the ratings, we normalized them per participant (by subtracting their minimum confidence rating and dividing by the difference between their maximum and minimum confidence values, like Charalampaki et al., 2023), to account for possible differences in the range of confidence ratings used by each participant. Further, in line with the recommendations, to avoid having empty bins that would hinder the fitting procedure (Maniscalco & Lau, 2012), when fitting the models for each participant, we added 1/(2*number of bins) = 1/12 to each input vector containing the count of each confidence rating bin type.

We used the “BayesFactor” package (Morey & Rouder, 2018) to obtain the Bayes factors in favor of H1 (BF_10_ values), the “rstatix” package (Kassambara, 2023) to estimate effect sizes and ran parametric correlations using the “ggstatsplot” package (Patil, 2021), all implemented in R (version 4.2.2, R Core Team, 2020). Prior to the statistical analyses, we log-transformed d’ and scaled (by mean-centering to the condition-specific mean, and normalizing by the condition-specific SD) the confidence ratings because they were not normally distributed. We report mean values and standard deviations (M = mean value ± SD).

### Electrophysiology (EEG) recordings

We recorded EEG data using 62 active electrodes fitted on an elastic cap following the 10–20 system g.tec position montage (g.tec medical engineering GmbH Austria). We placed two reference electrodes (one on each earlobe) and three electrodes around the right eye (above, below and outer canthus) to record horizontal and vertical eye movements. We used AFz as the ground electrode and the electrode on the right earlobe as the online reference. The signal was digitized at a sampling rate of 1200 Hz, online band-pass filtered between 0.5 and 200 Hz, and notch-filtered between 48 and 52 Hz to remove line interference. We monitored the electrodes’ impedances throughout the experiment to keep them below 10 KΩ.

### EEG analysis

We preprocessed the EEG signal using the EEGLAB toolbox (Delorme & Makeig, 2004) and custom scripts implemented in MATLAB (R2020a, MathWorks, Natick, MA). In line with our pre-registered plan, we re-referenced the EEG signal to the average signal of the earlobe electrodes, filtered the signal using separate finite high-pass (1 Hz passband edge) and used low-pass (100 Hz passband edge) Hamming windowed-sinc finite impulse response (FIR) filters, notch-filtered using a filter with passband edges at 49 and 50 Hz, downsampled to 600 Hz and run Independent Component Analysis (ICA; runica). We first flagged the components, then visually inspected them to reject those involving eye movements and muscle contraction (focusing on those with a probability of over 0.90 for belonging to the eye or muscle component category). After downsampling to 250 Hz, we then epoched the data time-locked to i) participants grabbing the ball (both for the intact and manipulated intervals, ii) participants releasing the ball (both for the intact and manipulated intervals), iii) the cue (i.e. the appearance on the screen of the agency question) for the discrimination response, and iv) the cue for their confidence rating. All epochs, save for the epochs based on the cue for confidence rating, started 1 s before the event of interest and ended 2 s after. They were baseline corrected for the time window -150 ms to 0 ms relative to each time-locked event. For the confidence cue epochs, the epoch started 1 s prior and ended 1 s after the cue. We first automatically rejected noisy epochs using the function *pop_autorej* with a threshold of 70 µV for extremely large fluctuations (Rejected epochs time-locked at: Release: M = 173.68 ± 77.85; Grab: M = 184.61 ± 82.87, Cue for discrimination: M = 102.87 ± 41.36; Cue for confidence: M = 107.11 ± 42.10) and then visually inspected the output to reject any remaining noisy epochs (Release epochs: M = 304.79 ± 107.47; Grab epochs: M = 284.92 ± 100.45, Discrimination cue epochs: M = 151.66 ± 55.39; Confidence cue epochs: M = 164.45 ± 56.10). We excluded the epochs time-locked to the discrimination cue from one participant because they were clearly noisy, but included all other epochs from this participant. We excluded the time-locked epochs at grabbing the ball from three participants because the time window between grabbing and releasing the ball was less than 400 ms for these three participants. Specifically, the time window between grabbing and releasing the ball for all 38 participants was 768 (± 204). Therefore, we included only trials in which the time window between grabbing and releasing the ball was at least 600 ms in both intervals. This process resulted in excluding epochs time-locked at grabbing the ball from 35 participants (92 ± 131) and all the epochs time-locked at grabbing the ball for three participants who did not have enough/or any trials surviving this exclusion procedure. Finally, we excluded a small set of epochs from those trials in which, due to a technical error, at least one of the triggers fired more than once, introducing uncertainty in the timing (Rejected epochs time-locked at: Release: M = 3.16 ± 2.95; Grab: M = 1.53 ± 2.35, Discrimination cue: M = 1.50 ± 1.46; no epochs were removed time-locked to the confidence cue). Therefore, for the final analyses we used the following number of epochs at each time-locking event: Grab: M = 394.29 ± 172.00.92; Release: M = 470.21 ± 143.54; Discrimination cue epochs: M = 234.32 ± 70.53; Confidence cue epochs: M = 224.97 ± 65.46. We chose to focus our analysis on the FCz electrode, based on previous studies on agency (comparing event related potentials related to primes, movement and outcome: Sidarus et at., 2017; ERP and time-frequency analysis: Wen et al., 2017) and scalp topography.

To compare the amplitude of the EEG signal for the epochs created, we fitted linear mixed-effect models using the "afex" package (Singmann et al., 2023). We then used cluster-based permutation tests to control for family-wise error rate (Maris & Oostenveld, 2007). For the latter, we permuted the factors’ labels (used on each model) on each trial and fitted linear mixed-effect models on each time-point 5000 times. For each main effect and interaction, we got clusters of neighboring time-points within the epoch when there was a significant difference between the conditions (p < 0.05). Then, we estimated the sum of the F-statistic for each cluster and saved the largest value. By repeating this process, we obtained the distribution of the sum statistics related to finding an effect by chance. We then tested whether the significant effects referred to time-points that were both lower than our alpha (0.05) and fell above the 95^th^ percentile of the time-points of the sum-statistics distribution.

For illustration purposes only, to explore how the the main effects of these factors and their interactions across the scalp, we fit the data from the remaining 61 channels using the same linear mixed effect models and then plotted the topographic maps of the mean beta estimates across the timepoints included in each of the significant clusters using the *eegUtils* package (Craddock, 2022) in R.

## Results

### Behavioral

First, we compared participants’ discrimination performance —measured with d’— and found that there was no significant difference between the two conditions (*d’_movement_* = 1.02 ± 0.11; *d’_outome_* = 1.04 ± 0.15; t(39) = -0.42, p = 0.677, Cohen’s d = -0.07, BF_10_ = 0.19; Fig. 2.A). That is, the manipulation did not significantly affect the discriminability of the intervals in terms of control. This result confirms that the staircased procedure employed to keep participants’ accuracy stable in the discrimination task worked, allowing us to validly compare participants’ Mratio values between the two conditions. Further, we found that mean confidence ratings did not differ between the two conditions (*Mean confidence_movement_* = 54.45 ± 15.03; *Mean confidence_outcome_* = 56.31 ± 13.51; t(39) = -1.71, p = 0.095, Cohen’s d = -0.27, BF_10_ = 0.64; Fig. 2.B). That is, we get weak evidence that participants’ confidence was comparable when they assess the accuracy of their representations of agency, both when the manipulation affected their movement or the outcome. Additionally, we found that participants’ metacognitive sensitivity, measured with metad’ was similar between the two conditions (meta*d’_movement_* = 0.86 ± 0.38; meta*d’_outcome_* = 0.86 ± 0.40; t(39) = 0.04, p = 0.968, Cohen’s d = 0.006, BF_10_ = 0.17; Fig. 2.C). Crucially, contrary to our pre-registered hypothesis, we found that metacognitive efficiency, measured with Mratio, did not differ between the two conditions (*Mratio_movement_* = 0.84 ± 0.36; *Mratio_outcome_* = 0.82 ± 0.34; t(39) = 0.25, p = 0.806, Cohen’s d = 0.04, BF_10_ = 0.18; Fig. 2.D). This shows that metacognitive representations of agency are equally sensitive to violations of predictions of the representations of the movement and outcome. This result held after removing two participants with negative Mratio (see Appendix).

**Fig. 2.**
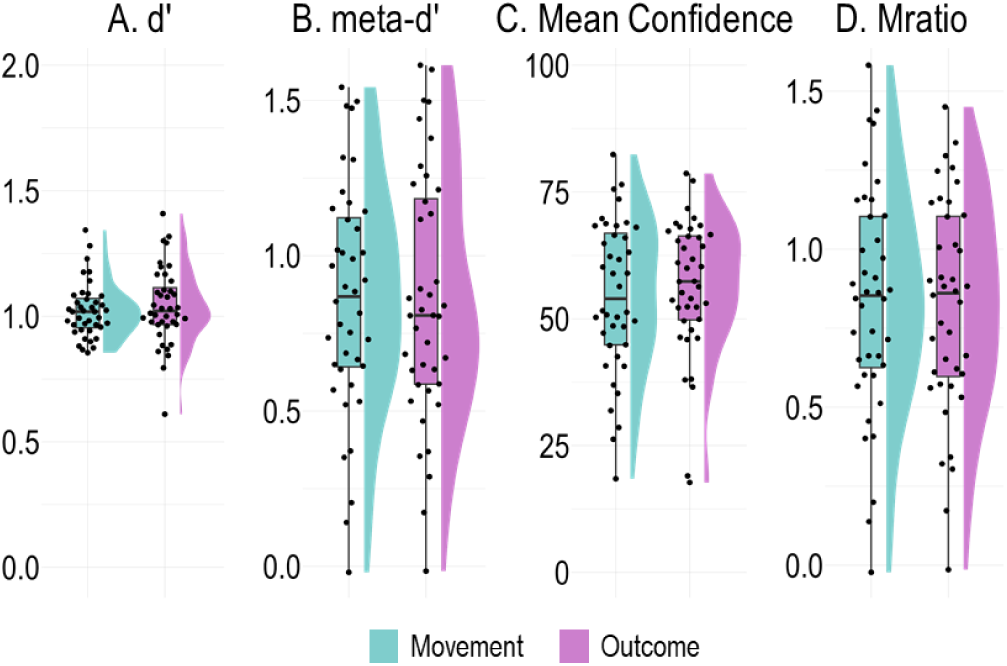
A. d’: Discrimination performance. B. metad’: Metacognitive sensitivity, C. Mean Confidence ratings and D. Mratio: Metacognitive efficiency. The violin plots depict the smoothed distributions of the data per condition and the boxplots depict the interquartile range. The black circles represent the estimates per participant

In an exploratory analysis, we then examined the relationship between metacognitive representations of agency in the two conditions. We wanted to test whether participants’ sensitivity to movement and outcome prediction violations were not only comparable but also correlated. We found that the correlation was not significant (t(38) = 0.36, Pearsons’s r = 0.06, p = 0.72, CI = [-0.26, 0.36], BF_10_ = 0.26, n = 40; Fig. 3). This result shows that although visual manipulations of the movement and the outcome might similarly affect metacognitive representations of agency judgments at the population level, they differentially do that for each participant.

**Fig. 3.**
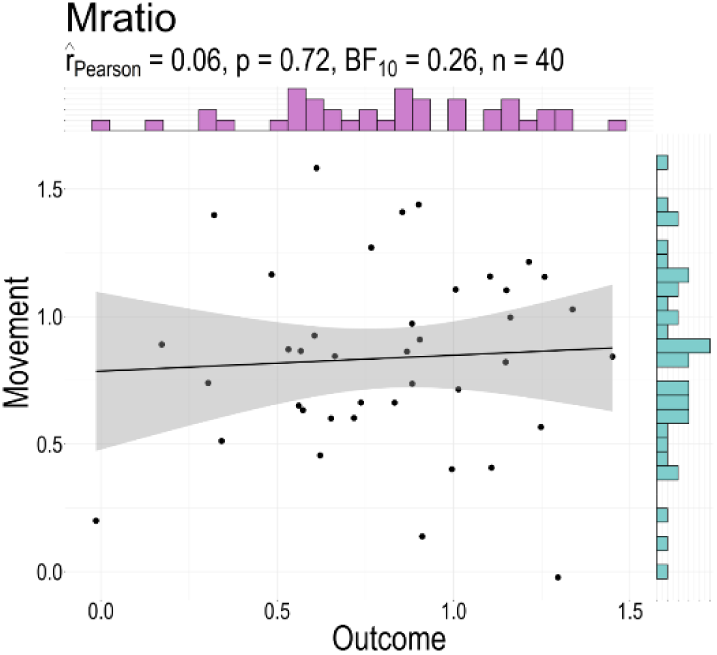
Correlation between the Mratios of the two conditions. No significant correlation was found between the Mratios of the *Movement* and *Outcome* condition. The black line represents the regression line and the shaded area illustrates the 95% confidence interval. The black circles represent the Mratio for each participant and the histograms the distribution of the data for each condition.

### Electrophysiology (EEG)

We then looked at the EEG signal to explore whether the different representation violations due to the movement or outcome manipulations are linked with distinct brain signals.

#### I. Movement epoch: arm movement

We first looked at the epochs time-locked at the point participants grabbed the ball. To compare the two manipulation conditions, we subtracted the amplitude of the EEG signal of the intact interval from that of the manipulated interval, separately for each condition. (We did this subtraction strictly for those trials where both epochs, intact and manipulated, could be included in the dataset). We expected that the signal would be different between the epochs only for the Movement trials, as these are the only trials in which we used this type of manipulation. We also expected that the amplitude difference would result from the interaction between the condition, participants’ accuracy in the discrimination response, and their confidence ratings indicating that whatever happens during these epochs informs both participants’ agency discrimination response and confidence ratings. Therefore, we tested our exploratory hypothesis using linear mixed effect models with the formula: EEG amplitude difference ∼ condition * confidence * accuracy in discrimination response + (1 | participant), which we tested for each time-point of our data. For this analysis, we focused on the time window from 0 to 600 ms after the participant grabbed the ball. After visually inspecting the movement traces from all the participants (hand position over time), we saw no behavioral indications that participants tested whether there was a movement manipulation within this time-window, as their movements had a consistent stereotypical outlook. In other words, they completed each throw in one pass. Cluster based permutation tests suggested the following clusters (we indicate each of their approximate onset times) to be larger than expected by chance: Three clusters linked to a three-way interaction between condition, accuracy and confidence (onset ∼ 240 ms, 344 ms and 468 ms), a cluster linked to the interaction between condition and confidence (onset ∼ 344 ms), two clusters linked to the interaction between confidence and accuracy (onset ∼ 236 ms and 428 ms), two clusters linked to the interaction between condition and accuracy (onset ∼ 268 ms and 348 ms), a cluster linked with a main effect of condition (onset ∼ 352 ms) and a cluster linked with a main effect of accuracy (onset ∼ 240 ms; Fig. 4). For illustration purposes, here and in the subsequent figures, we plot the topography, across the scalp, of the beta values for an effect identified in the timeseries. Here, the topoplot in the inset shows the beta estimates for the three-way interaction based on the condition, level of confidence, and accuracy averaged across the time window included in the shaded area.

**Fig. 4.**
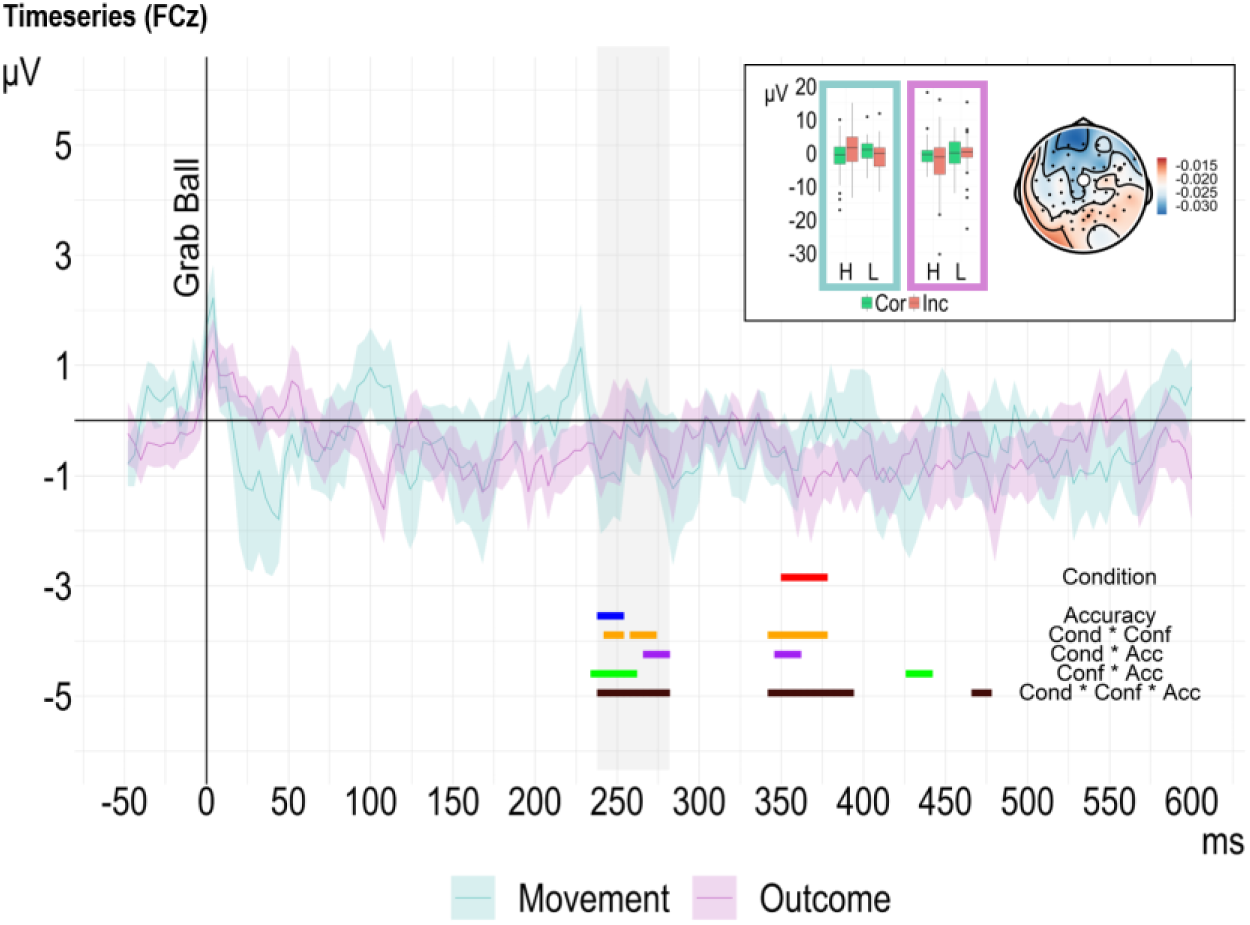
EEG amplitude difference between the manipulated and intact interval for the two conditions. The signal was time-locked to when participants grabbed the ball (FCz electrode). The significant time-point clusters from the cluster-based permutation analysis (EEG amplitude difference ∼ Condition * Accuracy * Confidence + (1|participant)) are color-coded to indicate the effect they represent, marked on the bottom-right. In the inset, the boxplots (left) show the mean EEG amplitude difference across the time points included in the first cluster linked with the three-way interaction (area marked with gray) based on the condition, level of confidence (H: high; L: low), and accuracy. The topographic map (right) shows the mean beta estimates across the timepoints included in the first cluster for all channels recorded. It indicates how the three-way interaction was related to the EEG amplitude in the remaining channels.

#### II. Outcome epoch: Ball flight

We followed a procedure similar to the one described above to compare the two conditions’ epochs when participants released the ball. That is, within each trial, we subtracted the EEG amplitude of the intact interval from the amplitude of the manipulated interval, respectively, and we then tested for significant differences with linear mixed effect models. We corrected for multiple comparisons using cluster-based permutation tests. We focused on the time window from 220 ms to 750 ms after participants released the ball for the difference between the epochs’ EEG amplitude. The reason was that always after participants released the ball, the ball disappeared from the screen for approximately 202 ms (M = 202.36 ± 0.21) because we wanted to avoid having participants identify the manipulation due to a visual jump of the ball’s position (see Methods). The permutation analysis revealed that there was a cluster linked to a three-way interaction between condition, accuracy and confidence (onset ∼ 280 ms) and two significant clusters due to the condition (onset ∼ 668 ms and 708 ms)

#### III. Report Cues

We then looked for differences in the EEG signal between conditions at the point participants were prompted to either report their discrimination decision or their confidence rating. For readability, from now on, we will call them “Cue1” and “Cue2” epochs respectively. To compare for differences among the time-points, we tested the following linear mixed effect model EEG amplitude ∼ condition * confidence * accuracy in discrimination response + (1 | participant) for each time-point of the Cue1 and Cue2 epochs. For both, we used a time window starting at 0 s and finishing at 500 ms from when participants were prompted. These time-windows were chosen based on the average time participants needed to respond (discrimination response: 651.36 ± 148.63 ms; confidence response: 844.10 ± 238.37 ms), to focus our analysis on the underlying signal linked with the discrimination task and avoid motor artifacts and other processes contaminate the neural signals.

Regarding the epochs time-locked to Response discrimination cue, the permutation analysis revealed that there were a few significant cluster linked to the condition (onset ∼ 12 ms, 104 ms, 248 ms and 372 ms), a cluster linked to confidence (starting around 256 ms), a cluster linked to the interaction between condition and accuracy (onset ∼ 356 ms) and a cluster based on the interaction between condition and confidence (onset ∼ 96 ms).The permutation analysis on the epoched data time-locked to Confidence cue, revealed a few significant clusters. The first one was linked to the factor condition (onset ∼ 24 ms), the second to the factor confidence (onset ∼ 136 ms) the third indicated an effect of accuracy (onset ∼ 36 ms) and finally a cluster indicating an interaction between confidence and accuracy (onset ∼ 40 ms).

## Discussion

We tested the role of movement and outcome information in metacognitive representations of agency. That is, we measured the clarity of the subjective experience of agency when either the visual feedback of a movement or its outcome are manipulated. Participants performed a goal-oriented action two consecutive times. Then, they discriminated on which of the two actions they felt more in control and rated their confidence in that decision. The advantage of this approach is twofold. First, with the discrimination agency task, we could objectively measure the accuracy of participants’ subjective sense of agency without biases (Wang et al., 2020). Second, by adding confidence ratings following the discrimination response, we could study the link between participants’ subjective confidence ratings and the accuracy of their agency discrimination responses. We quantified this link using the Mratio, which is, in principle, independent of first-order performance (Maniscalco & Lau, 2012, but see Guggenmos, 2021; Rausch et al., 2023). Therefore, we could directly compare movement and outcome representations, which cannot be otherwise compared as they differ on several dimensions (e.g., they happen at different time points during an action and they often have distinct qualitative characteristics). Overall, the results of the behavioral analysis and the exploratory analysis of the EEG signal are in line with the idea that movement and outcome prediction violations affect the experience of agency through different mechanisms. In what follows, we discuss each of these pieces of evidence separately and then bring them together to discuss how they might inform more theoretical accounts of agency.

First, the absence of differences in Mratio between the two kinds of manipulations suggests that metacognitive representations of agency are equally sensitive to movement and outcome prediction violations. This result goes against our pre-registered hypothesis that the mean Mratio would differ between the two conditions. Instead, we found in fact evidence for the null hypothesis of no differences between the conditions. Hence, these results suggest that the salience, or precision, of experiencing a violation of predictions following an action does not depend on whether those predictions are bodily or external. Because we measured the precision of metacognitive representations, and not, like previous literature, merely mean agency ratings, a direct comparison between these results and previous studies must be taken with caution. (These two representational levels —agency and metacognitive judgments about it— have been argued to be distinct, both behaviorally [Charalampaki et al., 2024] and computationally [Constant et al., 2022]). Nevertheless, these results do complement the field. Previous studies that addressed the relative importance of movement and outcome predictions’ violations on participants’ sense of agency have yielded conflicting findings. Some studies have argued that participants’ agency is more sensitive to outcome violations than movement violations (David et al., 2016; Oishi et al., 2018; Wen et al., 2015). Other studies have provided evidence for the opposite, namely that participants’ agency is more sensitive to movement violations relative to outcome violations (Metcalfe et al., 2013; Metcalfe & Greene, 2007a; Villa et al., 2018, 2021; Vogel et al., 2024). Typically, in this body of work, the manipulations compared are qualitatively different. For instance, in movement manipulations, the visual feedback is typically continuous and is presented for a sustained period of time; whereas in outcome manipulations the visual feedback is discrete, or for a shorter duration, and typically consists of a sound or color change at the end of the action (David et al., 2016; Wen et al., 2015; Oishi et al., 2018; Villa et al., 2018, 2021) or the outcome congruency (Villa et al., 2018, 2021; Vogel et al., 2024; Metcalfe et al., 2013; Metcalfe & Greene, 2007). This comparison is problematic because previous work has shown that the duration of the feedback can overshadow or conceal the true effect of movement and outcome prediction violations. Schmitter et al. (2021) found that visual feedback that was continuous differed from discrete feedback not only in participants’ accuracy in detecting a delay in the feedback, but also in the BOLD activity patterns. The authors interpreted these results as indicating that distinct predictive mechanisms underlie the processing of the two different types of manipulation (Schmitter et al., 2021). With these limitations in mind, in our study, we manipulated the visual feedback of both movement and the outcome continuously. Put simply, we manipulated either the visual representation of participants’ hand movement or the ball flight trajectory, which both have a long duration relative to the short-lived outcome representations employed in the previous studies. Therefore, we argue that the Skittles task allowed us to compare the two types of action representations devoid of response and comparison biases, in a tightly controlled fashion. This does not undermine the findings from previous studies but highlights the limitations of comparing simple subjective reports between experimental conditions that are known to participants.

Further, we found that participants’ Mratio following movement manipulations did not correlate with Mratio following outcome manipulations. Simply put, those participants in our sample who were most metacognitively sensitive to movement violations were not always those most sensitive to outcome violations. This result suggests that a simple comparison between mean judgments of agency following different kinds of manipulations might not be sufficient to reveal the dissociations between them: Two different manipulations might affect the experience of agency to the same extent and nevertheless be processed via distinct mechanisms. While our pre-registered hypotheses focussed on the mean differences between conditions, we nevertheless consider this result to suggest that the two kinds of violations are processed differently. One might wonder if goal-based expectations could explain this dissociation. Previous studies would support this interpretation. For example, Kumar and Srinivasan (2014) showed how goal-dependent the relative role of movement and outcome information is: in a multi-agent task, they found that when participants successfully achieved their goal, they rated agency independently of their motor performance. However, when participants missed their goal, they relied more on the sensorimotor information (Did what I did match what I saw?) rather than the outcome congruency to rate their agency. Further, van der Weiden et al. (2010) showed that there are inter-individual differences in the way participants represent their behavior (movement vs. outcome) and that these representations can be prompted by the use of subliminal primes (prior to the action) or specific instructions ("make a specific movement" vs. "reach a specific goal"). Together, these studies suggest, first, that reaching a goal can affect agency ratings following outcome manipulations differently than ratings following movement manipulations; and, second, that different participants can differ in these relative effects. Importantly, participants were always successful in meeting their goal in our study. That is, we set up the virtual Skittles scene in such a way as to ensure this was always the case. On rare occasions, when participants missed the target in either of the intervals, we excluded the trials from further analysis. Hence, we argue that this difference cannot explain our results.

Finally, our exploratory EEG analysis yielded some interesting findings that call for further testing. We contrasted the two manipulation conditions in two different ways. First, by comparing epochs while the manipulation was present (i.e., while participants were moving on *movement* trials —Fig. 4—, and while the ball was flying on *outcome* trials —Fig. 5) and second, by comparing epochs at the time of the first-order decision (Fig. 6). In the latter approach, the two manipulations are strictly comparable in both their visual input and motor output. We argue that, in the former approach, a direct comparison between EEG epochs is also granted, although they corresponded to different parts of the epoch and therefore to different visual input and motor output. Because we considered the difference signals between manipulated and intact intervals, this difference should reflect the salient mismatch signal from a comparison between expected and observed sensory consequences of an action. Arguably, then, this saliency signal is comparable between manipulation conditions.

**Fig. 5.**
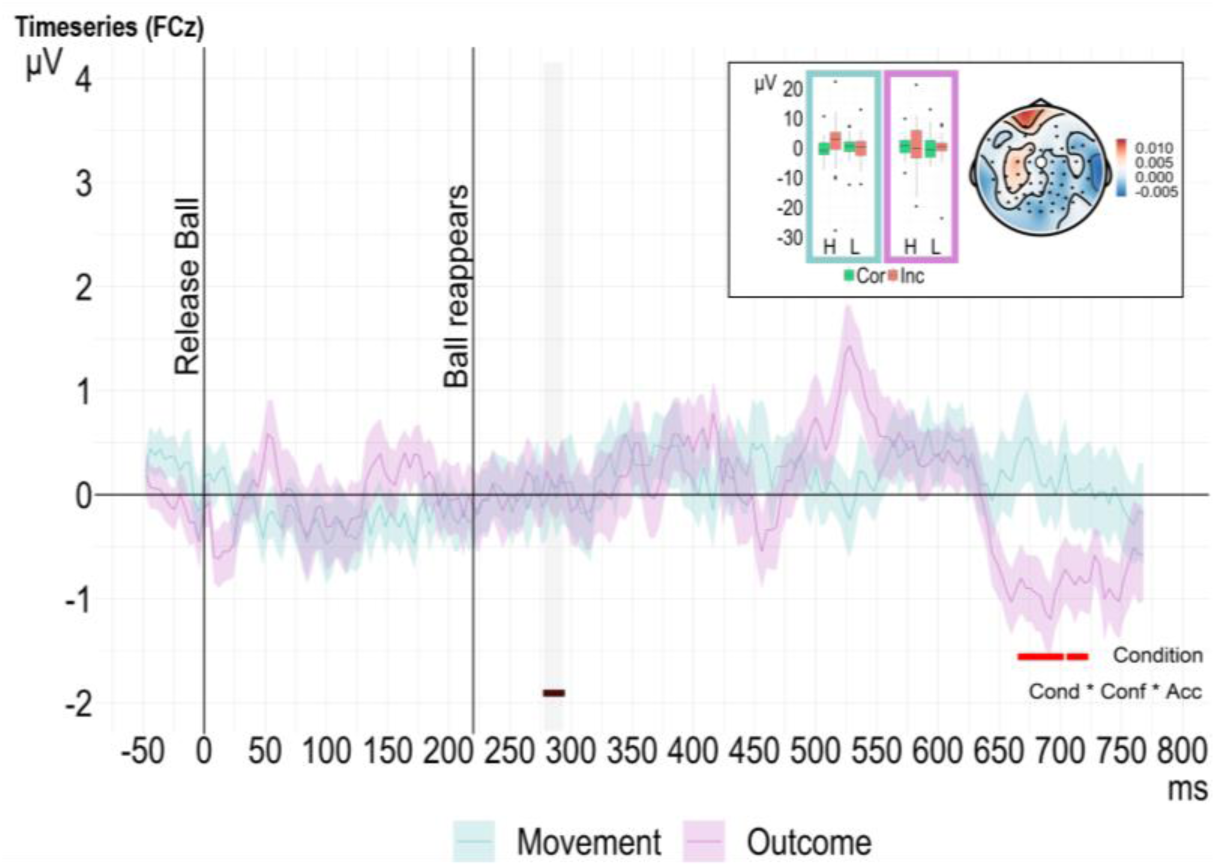
EEG amplitude difference between the manipulated and intact interval for the two conditions. The signal was time-locked to when participants released the ball (FCz electrode). Once participants released the ball it disappeared from the screen for 220 ms. The significant time-point clusters from the cluster-based permutation analysis (EEG amplitude difference ∼ Condition * Accuracy * Confidence + (1|participant)) are color-coded to indicate the effect they represent, marked on the bottom-right. In the inset the boxplots (left) show the mean EEG amplitude difference across the time points included in the first cluster which was linked with the three-way interaction (area marked with gray) based on the condition, level of confidence (H: high; L: low), and accuracy. As in Figure 4, the topographical map (right) shows the mean beta estimates across the timepoints included in the first cluster. It illustrates how the three-way interaction was related to the EEG amplitude difference in the remaining channels.

**Fig. 6.**
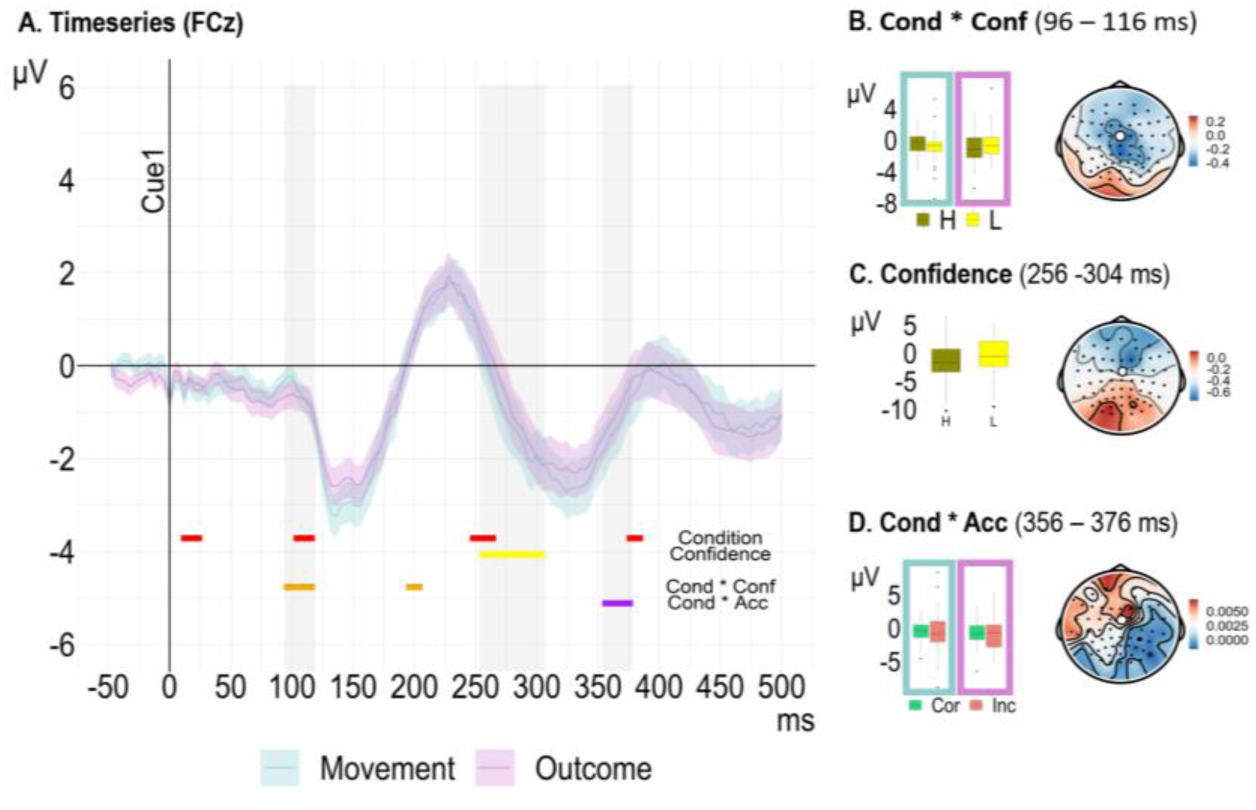
EEG amplitude for the two conditions (FCz electrode) time-locked to the onset of the discrimination cue (when participants provide their agency judgment). The significant time-point clusters from the cluster-based permutation analysis (EEG amplitude difference ∼ Condition * Accuracy * Confidence + (1|participant)) are color-coded to indicate the effect they represent, marked on the bottom-right (A.). On the right, the boxplots show the mean EEG amplitude across the time points included in the clusters (areas marked with gray) linked with the main effect of confidence (B.), the interaction between condition and confidence (C.) and the interaction between condition and accuracy (D.). The topographical maps (right) show the mean beta estimates across the timepoints included in the clusters of the EEG amplitude for each of the 62 channels.

When comparing the saliency signals from the two manipulation epochs, we found similar EEG correlates following the processing of violations of movement and outcome predictions: The EEG signal recorded during movement and outcome manipulations reflected both whether participants had correctly identified the manipulated interval and their confidence in that identification already within 250 ms after the manipulations took place (Fig. 4 and Fig. 5). However, only during movement manipulation did the EEG signal recorded reflect this relationship for a sustained interval (Fig. 4). In contrast, the EEG signal recorded during outcome manipulations did not vary with participants’ confidence, or with their discrimination accuracy (Fig. 5). In short, confidence had a measurable relationship with the EEG signal recorded during movement manipulations but not during outcome manipulations. To interpret this difference between conditions, we consider that confidence, like discrimination decisions, might rely on the saliency signals for prediction violations in the body. These might be automatically represented, or computed, with the reliability of bodily afferent signals. On the other hand, decisions about the effects of our actions in the environment might not be automatically computed and might correspond to a more reflective, post-hoc process (Haggard & Chambon, 2012).

Finally, in examining the EEG signal during the time of the discrimination decisions, we found it to reflect the accuracy and confidence of the decision along with the type of violation prediction. Together, the differences between the two manipulation epochs and the effects of condition at the time of the discrimination decision suggest that the underlying signal for agency reports differs between kinds of manipulation.

### Bodily and External representations and the underlying model of agency

To reiterate, our study showed that Mratios of the *Movement* and *Outcome* conditions did not correlate. We also found evidence for distinct EEG signals when we prompted participants to register their agency (discrimination) and confidence reports. These results cast doubt on a common underlying mechanism in processing violations of movement and outcome predictions in our sense of agency. The comparator model (Frith et al., 2000) is often used to explain the experience of (or reduction in) both the bodily and external sense of agency. There exists evidence to support the involvement of a comparator mechanism in the bodily sense of agency (but see Christensen & Grunbaum, 2018). However, the involvement of forward models in the comparison between predicted and perceived sensory consequences that go beyond the body has been challenged (Christensen & Grünbaum, 2018b; Dogge et al., 2019). While previous studies have shown that participants can predict the outcome of their actions in the environment before even receiving explicit feedback about it (Joch et al., 2017; Maurer et al., 2015), it cannot be automatically assumed that a copy of motor commands is responsible for the predictions about sensory consequences that go beyond the body (Christensen & Grünbaum, 2018b; Dogge et al., 2019). The comparator could explain the role of motor incongruencies and, by extension, bodily representation violations in a diminished sense of agency. If there were evidence that the comparator model could make predictions based on the forward models for sensory information external to the body, it would mean that the comparator could explain the role of outcome incongruencies and, by extension, external representation violations in a diminished sense of agency. However, based on our results, the same underlying mechanism seems less likely to be responsible for processing violations of both representations. A more inclusive model of agency, the Bayesian cue integration theory (Moore & Fletcher, 2012; Synofzik et al., 2013) posits that a sense of agency results from a weighted combination of different cues and puts forward a computational mechanism to integrate these cues, some of which result from a comparison, and others that may not. The weighting assigned to each of the cues can be determined by priors (prior knowledge and expectations). Because the model is, first, agnostic about how exactly these sources of information arise and, second, because it suggests that these cues are integrated, the cue integration model may be more suited to account for our results.

### Limitations

In our experiment we chose to study the role of bodily predictions violations by manipulating the visual representation of participants’ hand which was a rectangle displayed on the screen. One could argue that this is not a very accurate body part representation. However, we know participants do experience ownership over a variety of visual representations of their body parts (Kondo et al., 2018). Therefore, we consider our results still hold, but admittedly it would be interesting to replicate our results by comparing more bodily visual representations of participants’ movement.

### Future directions

We used a metacognitive task to study the subjective experience of agency devoid of response biases. Another option would be to study agency with implicit measures. For example, temporal binding consists of biases in the perceived timing of the movement and the outcome of voluntarily performed actions (Haggard et al., 2002): movement initiation is perceived as happening later, and outcome as happening earlier, than they really did. Alternatively, sensory attenuation, where the sensory consequences of voluntarily performed actions are perceived as less intense (Blakemore et al., 2000) compared to those of passive movements, is also sometimes used to quantify the sense of agency. However, these measures do not directly address the subjective component of agency; rather, they act as a proxy. In particular, there is an ongoing discussion on the link between implicit and explicit measures of agency, the details of which are outside the scope of this manuscript. Briefly, the discussion revolves around whether these two kinds of measures target the same phenomenal experience. While some studies argue that they do (e.g. Imaizumi & Tanno, 2019), others argue against it (Dewey & Knoblich, 2014; Schwarz et al., 2019). Therefore, it would be interesting to compare the role of bodily and external representations on agency with implicit agency measures. Not only to inform the discussion of implicit versus explicit measures, but also to compare our results using a metacognitive measure and those from implicit measures.

### Conclusions

We found that the sensory information of both the movement and the outcome is important for experiencing being the author of an action we intentionally perform. However, the underlying mechanisms that incorporate these two different sources of information into a comparison between expected and actual sensory feedback will likely differ. Further research is needed to understand how movement and outcome representations inform our sense of agency

## Data availability statement

This study was pre-registered (https://osf.io/sjurk), and any deviations from the pre-registered plan are explicitly reported. The preprocessed data, as well as R and MATLAB scripts for the analyses, are freely available at (https://osf.io/sm7rg/).

## CRediT authorship contribution statement

**Angeliki Charalampaki:** Conceptualization, Data curation, Formal analysis, Investigation, Methodology, Project administration, Software, Validation, Visualization, Writing – original draft, Writing – review & editing.

**Heiko Maurer:** Software, Writing -review & editing.

**Lisa K. Maurer:** Software, Writing - review & editing.

**Hermann Müller:** Software, Writing - review & editing.

**Elisa Filevich:** Conceptualization, Funding, Methodology, Project administration, Resources, Software, Supervision, Writing – original draft, Writing – review & editing.

## Acknowledgements

We thank Pedro Espinosa Mireles de Villafranca and Paula Guiomar Alarcón de Antón for support with experiment setup and data collection.

## Appendix

### Excluding participants with negative Mratio

Two of the participants had negative Mratio. Therefore, we ran all the analyses excluding these two subjects to test if the results hold. In line with the results we report in the main text, we found no significant difference between participants discrimination performance (d’) in the two conditions (*d’_movement_* = 1.03 ± 0.02; *d’_outcome_* = 1.04 ± 0.03; t(37) = -0.23, p = 0.82, Cohen’s d = -0.04, BF_10_ = 0.18). Further, there was no difference in participants confidence rating (*Mean confidence _movement_* = 54.67 ± 2.40; *Mean confidence _outcome_* = 56.21 ± 2.23; t(37) = -1.47, p = 0.151, Cohen’s d = -0.24, BF_10_ = 0.47) or metacognitive sensitivity (meta*d’_movement_* = 0.90 ± 0.05; meta*d’_outcome_* = 0.87 ± 0.06; t(37) = 0.40, p = 0.694, Cohen’s d = 0.06, BF_10_ = 0.19). Lastly, similar to the results we report in the main text participants Mratio was not different between the two conditions (*Mratio_movement_* = 0.88 ± 0.05; *Mratio_outcome_* = 0.83 ± 0.05; t(37) = 0.70, p = 0.497, Cohen’s d = 0.11, BF_10_ = 0.22).

Excluding the two participants from the correlation analysis had no effect on the correlation results (t(36) = 0.24, Pearsons’s r = 0.04, p = 0.81, CI = [-0.28, 0.35], BF_10_ = 0.25, n = 38). Thus, the absence of correlation between the Mratios of the two conditions we report in the main text holds.

